# Monovalent Cation Dependent Oligomerization and DNA Binding of MoyR: a GntR family monooxygenase regulator in *Mycobacterium tuberculosis*

**DOI:** 10.1101/2025.08.21.671446

**Authors:** Thanusha D. Abeywickrama, Hideaki Tanaka, Genji Kurisu, Inoka C. Perera

**Affiliations:** Synthetic Biology Laboratory, Department of Zoology and Environment Sciences, Faculty of Science, University of Colombo, Colombo 00300, Sri Lanka; Laboratory of Protein Crystallography, Institute for Protein Research, Osaka University, 3-2 Yamadaoka, Suita, Osaka, Japan; Current address: Section for Plant Biochemistry, Department of Plant and Environmental Sciences, University of Copenhagen, Thorvaldsensvej 40, 1871 Frederiksberg; Current address: TechnoPro, Inc., TechnoPro R&D Company, Hyogo 650-0047, Japan

**Author notes:** Correspondence: Inoka C. Perera, **Email:**.

**Keywords:** Antibiotic resistance, HutC family regulator, Oligomerization, DNA-binding, Tetrameric protein

## Abstract

Bacteria constitute a large domain of prokaryotic microorganisms with a remarkable ability to perceive environmental stimuli, and respond efficiently to environmental stresses by regulation of gene expression mediated by transcriptional regulators. *M. tuberculosis* is a classic example of this, where ample studies have provided evidence of how the transcriptional regulators are involved in various survival mechanisms in the bacterium. *Mycobacterium tuberculosis* is a well-known pathogen due to the emergence of drug resistance associated with it, and the genome of *M. tuberculosis* encodes many transcriptional regulators. Our work explicates the mechanism of K^+^/ Na^+^ mediated oligomerization and DNA binding of MoyR. *In vitro* assays revealed a complex pattern of MoyR oligomerization depending on monovalent cation concentrations where MoyR tends to form tetramers and hexamers which directly affect the MoyR affinity to its cognate DNA. The tetramer bound DNA was found to be highly stable even at high monovalent cation concentrations such as 500 mM which is unusual for a protein-DNA binding in a mesophilic bacteria. GntR proteins are substantially known to bind the DNA as homodimers in a two-fold symmetry, but this study is the second, that reports of a tetrameric GntR protein assembly after the GntR regulator Atu1419 in *Agrobacterium fabrum*. MoyR binds with high affinity (K_d,app_ = 3.47 nM) and specificity to the shared promoter region between divergently oriented Rv0789c-Rv0790c-Rv0791c-*moyR* operon and Rv0793. The *moyR* gene cluster was identified as a conserved region and most of the adjacent genes are homologous to monooxygenases which highly likely to be likely to be inplicated in a polyketide antibiotic synthesis pathway of the bacterium and this can provide a novel paradigm to the antibiotic tolerance and resistance of *M. tuberculosis* wherein this elicits the importance of MoyR as a drug target.

## Introduction

*Mycobacterium tuberculosis* included in the *Mycobacterium* genera is known simply as the causative agent of the disease tuberculosis, claiming millions of lives each year over the course of human history. Being one of the major antibiotic-resistant respiratory pathogens, *M*. *tuberculosis* possesses a naturally high resistance level to antibiotics, while newly acquired mutations confer further resistance. As a microorganism associating humans for years in recorded history, it poses various mechanisms to vanquish the stresses and adapt to survive in the human host. Efficient regulatory mechanisms that control the expression of gene products that are necessary to mount the survival response and virulence of the bacterium underlines its success as a pathogen. Hence, transcriptional regulators are considered as the key molecules that gives rise to a tight regulation of genes in cellular pathways during infection, colonization, antibiotic-resistant, virulence and persistence inside the hosts.

Genome of *M*. t*uberculosis* encodes more than one hundred putative transcriptional regulators. Out of which, GntR family of transcription factors are found associated with cellular metabolism or responding to changes in the immediate environment. First described by Fujita *et al*. in *Bacillus subtilis* (Fujita et al. 1986), these GntR regulators are comprised of two domains N and C-terminal domains, where the N-terminal DNA-binding domain is remarkably conserved within the GntR family by having a winged helix-turn-helix motif (wHTH) (Aravind *et al*., 2005; Gajiwala & Burley, 2000; Rigali *et al*., 2002). Comparatively, a high structural heterogeneity can be found in the C-terminal ligand-binding domains within the family, accordingly, GntR family is subdivided into seven subfamilies FadR, HutC, MocR, YtrA, AraA, PlmA and DevA based on the findings by Rigali *et al*., Lee *et al*., Hoskisson *et al*. and Franco *et al*. (Franco *et al*., 2006; Hoskisson *et al*., 2006; Lee *et al*., 2003; Rigali *et al*., 2002). The C-terminal domain of the GntR regulators consists of a binding site that can accommodate a small effector molecule. Binding of the ligand triggers a conformational change that regulates the affinity of the N-terminal DNA-binding domain for the operator sequence, thereby regulating the expression of adjacent genes (Wriggers, Chakravarty, and Jennings 2005).

Vindal *et al*. reported 07 putative GntR regulators in the genome of *M*. *tuberculosis* and five out of seven classified as FadR regulators, Rv0043c, Rv0165c, Rv0494, Rv0586 and Rv3060c (I and II), whereas Rv0792c and Rv1152 classified as HutC and TyrA-like protein respectively (Vindal, Ranjan, and Ranjan 2007). Out of these regulators, many are uncharacterized to date whereas only two of the regulators Rv0494c and Rv1152 were identified as a fatty acid-responsive transcriptional regulator and an antibiotic resistance regulator respectively (Biswas et al. 2013) (Zeng et al. 2016). These findings indicate that the GntR regulators in *M*. *tuberculosis* could play a major role in pathogenicity and further signifies the importance of characterizing them in order to unveil the underlying mechanisms with the aspects of identifying novel drug leads in order to overcome the problem of antibiotic resistance. The regulator Rv0792c in the HutC subfamily was recently identified as a monooxygenase regulator in *M. tuberculosis*, named as MoyR which can be involved in a polyketide metabolism pathway of the bacteria (Abeywickrama and Perera 2021). Herein we report the behavior of MoyR in the presence of monovalent cations and its DNA binding mechanism which is important for its repressor functions. Mechanistic data reported here is crucial for the development of therapeutic agents against tuberculosis to battle drug resistance.

## Materials and Methods

### Cloning, Protein Expression and Purification

The gene encoding MoyR was amplified by PCR from *M. tuberculosis* H37Rv genomic DNA using forward primer 5′-CTT **GAA TTC** ATG ACA TCT GTC AAG CTG GA - 3′ and reverse primer 5′-CGT **GAA TTC** TCA TGC GAA ATC TCG TTT CT - 3′ which introduces a EcoR1 recognition sequence (in boldface) to both the primers. The PCR amplicon was cloned into the expression vector pET28-a(+) (Novagen^®^) which carries an N-terminal-6X histidine fusion tag and transformed into *E. coli* DH5-α cells. The correct construct was confirmed by sequencing. The verified construct was then transformed into *E. coli* BL21(DE3)pLysS (Novagen^®^) cells for overexpression. A well isolated single colony was picked from a freshly streak plate to inoculate an overnight culture. This overnight culture was grown in Luria-Bertani (LB) broth of pH 7.0 with 50 µg/ml kanamycin at 37℃ with a shaking speed of 190 rpm. The starter culture was diluted in 1:500 in LB broth of pH 7.0 with (50 µg/ml kanamycin) (2 L total volume). Cultures were grown at 37 ℃ (200 rpm) and protein expression was induced with 0.5 mM Isopropyl β-D-1-thiogalactopyranoside (IPTG) at an optical density of ⁓ 0.5 at 600 nm. Induced culture was grown for 3 hours, chilled on ice for 30 min and cell pellets were harvested by centrifugation and stored at-80 ℃. The cell pellet was resuspended in 5 volumes of ice-cold lysis buffer [50 mM Potassium phosphate buffer (pH 7.6), 150 mM KCl, 5% Glycerol, 10 mM Imidazole, 0.15 mM phenylmethylsulfonyl fluoride (PMSF), 10 mM 2-mercaptoethanol] and treated with 200 µg/mL lysozyme for 1 hour under chilled conditions. Lysis was completed with 0.05% Triton X-100 and immediate addition of 500 mM NaCl to the suspension. The cell suspension was sonicated for 10 min under chilled conditions and DNase I was added to digest nucleic acids. This suspension was centrifuged under a high centrifugal force. The supernatant was loaded on to a Ni-NTA agarose (1 mL/ min) column which was pre-washed and equilibrated with the lysis buffer. The column was washed with 5 bed volumes of wash buffer [50 mM Potassium phosphate buffer (pH 7.6), 150 mM KCl, 20 mM Imidazole] and protein was eluted with a linear imidazole concentration gradient from 10 mM to 250 mM. A sample of each of the flow through and eluted fractions were subject to SDS-PAGE to identify the purest fractions and the selected purest fractions were pooled and dialyzed for 3 hours against dialysis buffer [20 mM Potassium phosphate buffer (pH 7.0), 150 mM KCl, 5% Glycerol, 20 mM EDTA, 0.15 mM PMSF, 10 mM 2-mercaptoethanol]. The glycerol content of the protein sample was increased up to 20% for storage at-80℃. The 6X-histidine tag of the purified MoyR protein was cleaved with thrombin where 50 U/ mg of thrombin was added to 800 µg of protein followed by incubation with Ni-NTA agarose at 20 ℃ overnight. Undigested protein and cleaved poly-histidine were removed by centrifugation and identified with SDS-PAGE. Purity of the pooled fraction was determined by SDS-PAGE where the concentration of purified protein was measured by the BCA protein assay kit (Thermo Scientific) using bovine serum albumin (BSA) as a standard.

### Oligomeric states of MoyR Gel Filtration

To determine the oligomeric state of MoyR, gel filtration chromatography was carried out in two different salt concentrations, 150 mM and 500 mM. MoyR protein sample was equilibrated in ice for 1 hour and then filtered through 0.22 µm filter (Millipore) to remove any particles and the sample volume was adjusted to 5 ml. The diluted protein was injected to a HiLoad Sperdex^TM^ – 75 column (2.6×60 cm) (HiLoad 16/600, GE Healthcare) which was pre-equilibrated PBS with the desired salt concentration and eluted with the same buffer as the mobile phase buffer with a flow rate of 0.5 ml min^-1^. Elution profiles were monitored at 280 nm and the molecular weight of eluted proteins were estimated using a calibration standard using five homologous series of globular protein size markers Thyroglobulin (670 kDa), gamma-Globulin (158 kDa), BSA (67 kDa), Ovalbumin (44 kDa) and Myoglobulin (17 kDa). The gel phase distribution coefficient (K_av_) for each standard and MoyR was calculated using the equation 𝐾_𝑎𝑣_ = (𝑉_𝐸_ − 𝑉_𝑂_)⁄(𝑉_𝑇_ − 𝑉_𝑂_) where *V_E_* – elution volume of the protein *V_O_* – void volume of the column and *V_T_* – total column volume. The molecular weight (M_r_) of MoyR was determined from the standard curve as a plot of K_AV_ versus log M_r_ (Ó’Fágáin, Cummins, and O’Connor 2017).

### Glutaraldehyde Mediated Cross-linking

*In vitro* crosslinking of MoyR was mediated in reactions containing freshly prepared 0.1% glutaraldehyde, 14 µM MoyR in 50 mM sodium phosphate buffer (pH 7.5) and increasing concentrations of *moyO*-119bp (0, 1, 2, 3,4 µM) in a total volume of 20 µl. Reactions were incubated at 37 ℃ for 30 min and then they were terminated by adding 1 M Tris HCl at pH 8.0. The samples were electrophoresed at 150 V on a 15% SDS-PAGE gel. Molecular weights of covalently bound oligomers were determined using a broad range protein marker and its migration distance on the SDS-PAGE.

### DNA binding dynamics of MoyR

To identify the binding motifs of MoyR, the upstream region of *moyR* was analysed based on the reported HutC consensus binding motif signature and were analysed using MEME suite 5.2.0 with a given minimum of 10-bp length (Bailey et al. 2009). Identified putative HutC orthologs were aligned using ClustalW and consensus sequence logo was generated with WebLogo tool (Crooks et al. 2004).. The intergenic region *moyO* (119-bp) with the predicted binding sites was PCR amplified using forward primer 5′-AGGTCCAGCTTGACAGATGTCAT-3′ and reverse primer 5′-TACTGGCGCTG CCCACTGC AGTA-3′ and the PCR product was purified by electro-elution. To analyze the binding of MoyR with *moyO*, electrophoretic mobility shift assays (EMSA) were carried out as described previously with minor modifications (Perera and Grove 2010; Hellman and Fried 2007). A 10 µl reaction containing 3 nM *moyO* DNA, 1X EMSA binding buffer [10 mM Tris-HCl (pH 8.0), 100 mM KCl, 1 mM EDTA, 0.5% Triton X-100, 0.02 mg/ml BSA, 5% Glycerol] with MoyR protein in increasing concentrations was incubated at 25 ℃ for 30 min. Binding assays were carried out in triplicates. Then the reaction mixtures were electrophoresed using 1X TAE buffer on 8% polyacrylamide gels (39:1 [w/w] acrylamide: bis-acrylamide) at a voltage of 9 Vcm^-1^ for 1.5 hour at 4 ℃. The complexes and free DNA were quantified using ImageJ software (Perez and Pascau 2013). To determine the K_d_ of protein-DNA binding, the percentage of complex formation was plotted against [MoyR] and GraphPad Prism^®^ (GraphPad software) was used for fitting the data to Y = B_max_ x [MoyR]^n^⁄ K_d_ + [MoyR]^n^ equation where Y-fractional saturation, B_max_ – maximum specific binding, n – Hill coefficient, and K_d –_ apparent dissociation constant in the equilibrium (Ranganathan et al. 2018). To identify the MoyR binding motifs within the *moyO* intergenic region, EMSAs were carried out with short DNA oligonucleotides consisting specific binding motif and nonspecific regions as well. Customized double stranded DNA fragments were synthesized by BEX Co., Ltd, Japan. An amount of 0.32 pmol from the DNA fragment (*moyO*f_1_/ *moyO*f_2_/ *moyO*f_3_/ *moyO*f_4_/ *moyO*f_5_/ *moyO*f_6_) was incubated at 25 ℃ with increasing concentrations of MoyR as described above (Figure 4a). For each of the oligonucleotides EMSAs were carried out separately as mentioned and K_d_ was calculated. For the assay of stoichiometry 1.3 pmol of 119-bp *moyO* was incubated with increasing concentrations of MoyR in 1X EMSA binding buffer in a final volume of 10 µl at 25 ℃ for 30 min. The complexes were resolved by vertical electrophoresis in a 8% polyacrylamide gel in 1X TAE buffer at 9 Vcm^-1^ for 1.5 hour at 4℃ after a pre-run for 30 min. Gels were soaked in 0.5 µg/ ml ethidium bromide solution and complexes were visualized under UV light. The percentage of complex formation was plotted against [MoyR]/ [*moyO*] and the stoichiometry of complex formation was extrapolated algebraically *via* the X value at the intersection between tangents to the linear portion of the graph. The specificity of the binding was assessed by a competition assay in which 3 nM of *moyO* was challenged with increasing concentrations of non-specific linear pET28a+ plasmid DNA where the MoyR concentration kept constant based on the half maximal saturation value of the protein. Then the EMSA was carried out as described.

### Monovalent cation and pH Dependent DNA Binding

In order to determine the monovalent salt dependent MoyR-*moyO* complex formation, 3 nM of the *moyO* was incubated with 12 nM MoyR in a binding buffer (20 mM Tris-HCl, 1 mM EDTA, 0.5% Triton X-100, 0.02 mg/ml BSA, 5% Glycerol). Increasing concentrations of KCl or NaCl (0 to 500 mM) were added to a final volume of 20 µl and incubated the reactions at 25 ℃ for 30 min and electrophoretic mobility shift assay was carried out as mentioned before. Divalent cation dependency was also determined as in the same manner described above with increasing concentrations of MgCl_2_ or CaCl_2_. To determine the effect of pH for MoyR-*moyO* binding, three buffers were used in the binding reactions. The value pH 8.0 was maintained using Tris-HCl buffer, pH 7.0 was maintained using HEPES [(4-(2-hydroxyethyl)-1-piperazineethanesulfonic acid)], pH 6.0 and pH 5.5 was obtained using MES [2-(N-morpholino) ethanesulfonic acid] buffer. An amount of 3 nM *moyO* was incubated with increasing concentrations of MoyR in the desired binding buffer while the KCl concentration was kept constant at 100 mM. The incubation reactions and electrophoresis were carried out as described. The K_d_ values for each pH were determined accordingly.

### Protonation-Deprotonation and Particle Size Analysis of MoyR

In relation to the pH responsive binding of MoyR, in silico analysis of the monomeric MoyR was carried out to discern the protonation-deprotonation of the possible titratable residues using H++ server and PlayMolecule server (Gordon et al., 2005; Martinez-rosell et al., 2017). Previously identified conserved residues of MoyR were analyzed for protonation and the pKa values of the residues were determined to identify the pH sensitive critical residues that can participate in pH sensitivity. In the H++ server dimeric MoyR PDB structure was used for the analysis wherein the desired salt concentration was set to 0.15 M and the internal dielectric value was set to 10 or 6 according to the surface and buried residues. The pKa values of the titratable residues were determined at pH 8.0, 7.0, 6.0 and 5.0. The PlayMolecule server is also used to analyze the protonation of the residues. To estimate the particle sizes upon pH changes, a dynamic light scattering was carried out. Purified 2 mg/ml MoyR protein was dialyzed against different pH buffers at 4℃ for overnight to exchange MoyR into different buffers, Tris-HCl 7.5, HEPES 7.0, MES 6.5, MES 6.0 and MES 5.5. Followed by the dialysis, the MoyR protein buffers were filtered on 0.02 µm inorganic membrane AnotopTM 10 Plus filters from Whatman. A volume of 10 µl from the filtered protein samples were carefully pipetted into a disposable cuvette and slightly centrifuged to eliminate the air bubbles. Measurements were taken in replicates in a 658 nm light beam at 20℃ with the use of DynaPro® NanoStar® Dynamic Light Scattering Detector (Wyatt Technology Cooperation).

### Virtual Screening Analysis

The modelled structure of MoyR, Rv0791 and Rv0793 were used for molecular docking to determine the binding of different possible natural ligands using AutoDock 4.2.6 software (Morris, 2010). Other modelled structures were eliminated due to their low reliability of the models. Rigid-flexible docking was carried out where the protein molecule was set to a rigid file and the ligand was flexible. Genetic algorithm was used as search parameters and Lamarckian Genetic Algorithm was used as docking parameters. Virtual screening platform was established using PyRx software (Dallakyan & Olson, 2015). A rigid protein file in pdbqt file format and ligands were fed in SDF file format. Energy minimization was done for each ligand using uff (United Force Field) as the force field and conjugate gradients was used as the optimization algorithm with limitation of 200 iterations for each ligand. More than 400 possible natural ligands were extracted from the KEGG PATHWAY database. Ligands with lowest binding affinities (kcal/ mol) were selected and the binding interactions were analyzed using Discovery Studio Visualizer (Accelrys Discovery Studio Visualizer v 3.5. San Diego: Accelrys Software Inc, San Diego: 2010.) and UCSF Chimera (Pettersen et al., 2004) to identify the possible natural ligands for in vitro assays.

## Results

### Oligomeric States of MoyR

To determine the oligomeric state of purified MoyR, gel filtration chromatography was used MoyR protein in two different salt concentrations was subjected to gel filtration and eluted. The elution profile of MoyR with 150 mM NaCl showed as a single peak in the chromatogram which indicates a single species (Figure 1A). The elution profile of MoyR with 500 mM NaCl showed as one major peak with two other minor peaks which is indicative of multiple states of MoyR (Figure 1B). Theoretical monomeric weight of MoyR+His tag is 32.4 kDa and calculated molar weight of MoyR from calibration curve was 63.9 kDa which indicates a dimer. The other two molar weights were 139.9 kDa and 223.8 kDa which are correspond to a tetramer and a hexamer respectively (Figure 1C). Crosslinking of MoyR was mediated by adding 0.1% glutaraldehyde to determine covalently linked oligomeric structures and the molecular weight was determined according to the migration of the oligomers (Figure 1D). According to the interpolated values from the curve, the observed bands corresponded to MoyR monomer, dimer and hexamer weights. A faint band was visualized between the dimer and hexamer which could possibly be the tetramer.

**Figure 1:**
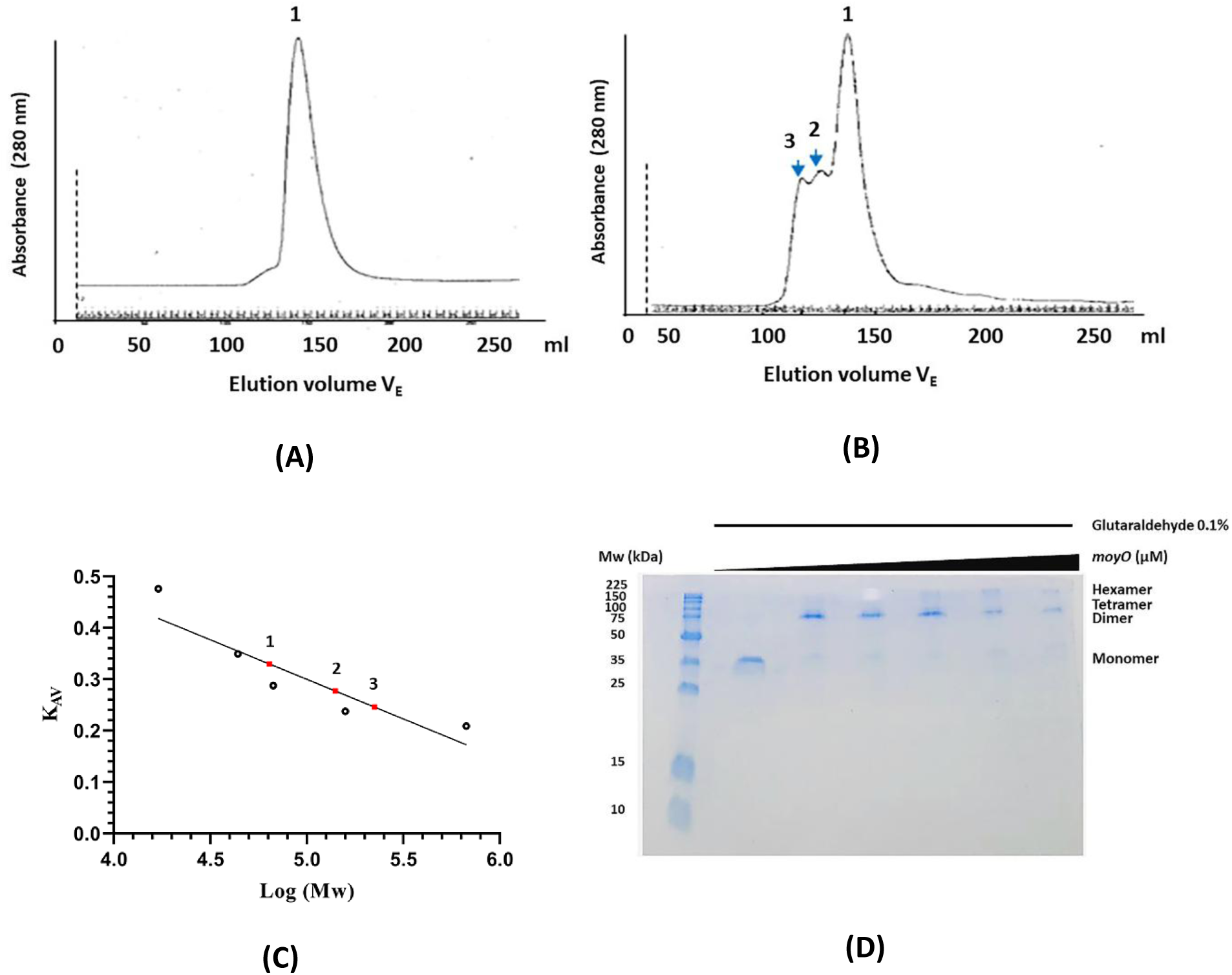
Oligomerization of MoyR. (A) Absorbance of MoyR PBS with 150 mM NaCl which is indicative of a single state of MoyR. The peaks indicate the absorbance at 280 nm in a particular elution volume and the elution volume varies with the molecular weight of the eluted species. (B) Absorbance of MoyR eluted in PBS with 500 mM NaCl which is indicative of multiple states of MoyR. (C) Plot of K_AV_ vs log molecular weight used to determine the molecular weight of MoyR and oligomers. Molecular weights of MoyR states are, 1 - 63.9 kDa, 2 – 139.9 kDa, and 3 – 223.8 kDa. (D) SDS-PAGE of glutaraldehyde mediated covalent cross-linking of MoyR. Lane 1-molecular weight marker, lane 2 – reaction without glutaraldehyde, lane 3 – 7 with increasing concentrations of binding site *moyO*.

### Operator Sites within the Upstream Region of moyR gene

Many HutC regulators are organized in two divergently transcribed operons, and this was observed in the *moyR* gene locus where the intergenic region was shared by Rv0789c-Rv0790-Rv0791c-*moyR* operon and Rv0793 (Figure 2A). As described previously *moyR* genomic locus consist of many hypothetical genes which are predicted to be genes encoding a group of monooxygenases (Abeywickrama and Perera 2021). The intergenic region between *moyR* and Rv0793 was mapped, and potential binding motifs were identified. Consensus sequence for the HutC family is NyGTMTAKACNy where the centre of the palindrome is usually conserved while the peripheral nucleotides vary (Sébastien Rigali et al. 2002; S. Rigali 2004). One binding motif with HutC signature was identified in the intergenic region of *moyR*-Rv0793 which is highly similar to the binding motif of fatty acyl responsive regulator FarR in *E. coli*. Binding motif of FarR has been experimentally determined and the identified that the primary site is located in a 21-bp region containing two 10-bp consensus sequences, TGTATTAA/TTT which are imperfect direct repeats.

**Figure 2:**
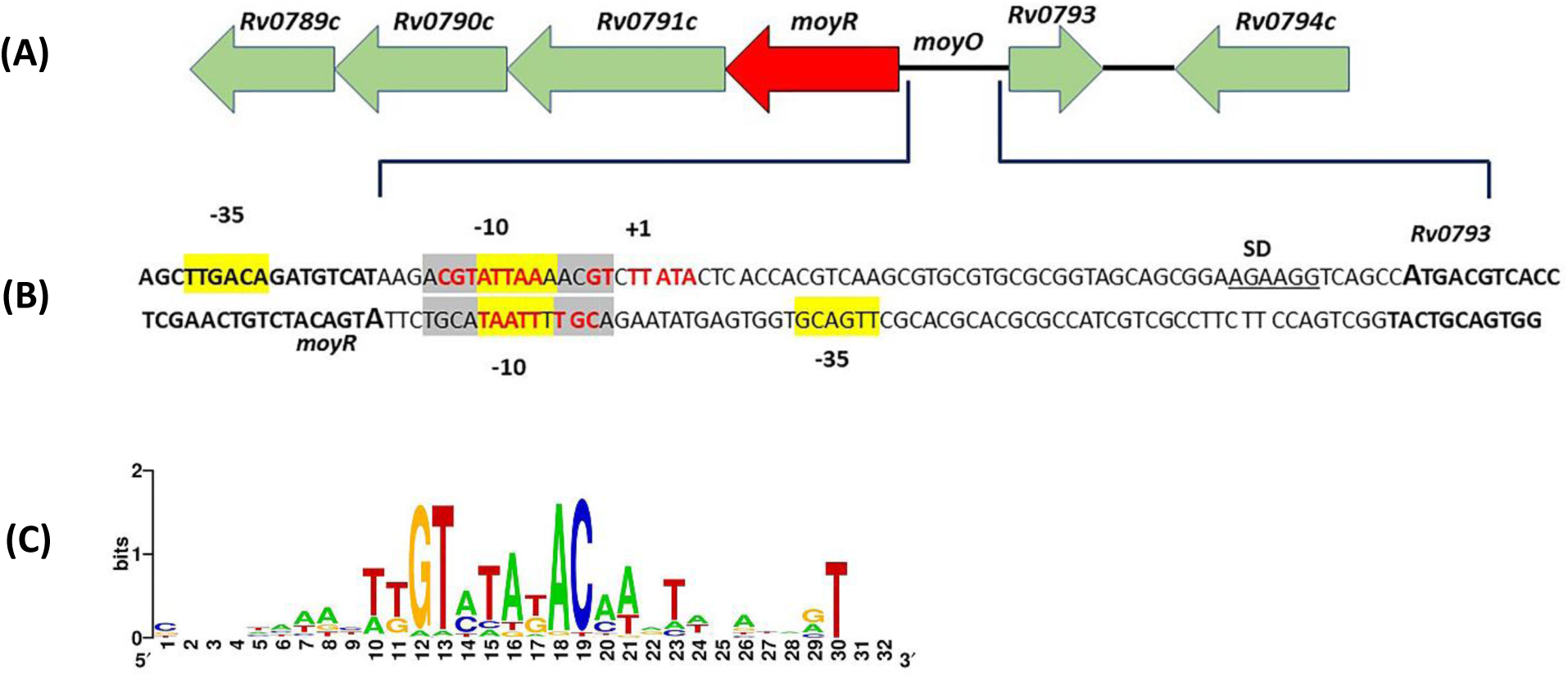
Operator site of MoyR. (A) Gene locus of *moyR*, 72-bp intergenic region located in divergently transcribed Rv0790c-Rv0791c-*moyR* and Rv0793. (B) Promoter region of MoyR. Predicted transcription start site indicated in +1, where the-10 and-35 hexamers are highlighted in yellow. The MoyR binding half sites are shown in bold arrows. The conserved base pairs of MoyR binding site compared with FarR binding site is highlighted in red. Short palindromic sequences are highlighted in grey. (C) Sequence conservation of MoyR binding site and sequence logo was generated using Weblogo server.

This pattern was highly conserved in the identified MoyR binding motif in the *moyO* region as 8-bp imperfect direct repeats which were able to identify only on one site within the intergenic region. Accordingly, MoyR operator sequence can be given as a 18-bp region containing two copies of 8-bp imperfect direct repeat, ACGTATTA separated by 2 base pairs (Figure 2B). The two homologous of MoyR, naming DasR and NagR recognize a highly similar operator site called *dre*-site which is termed after DasR responsive elements from *S. coelicolor* (Fillenberg et al. 2015). The *dre* consensus sequence ACTG**GT**CTAC **AC**CATT can be found in upstream regions of many genes in the genome of *S. coelicolor* where DasR plays a global regulatory role (Colson et al. 2006). A Weblogo analysis was carried out using selected HutC regulators (Suvorova, Korostelev, and Gelfand 2015) to identify the binding motif sequence with aid of experimentally determined consensus as well. HutC motifs revealed a highly conserved GT and AC region with a distance of 4 nt. (Figure 2C). The distance between the GT and AC of the consensus binding motif of *moyR* is 5/6 nt. To identify the motif occurrence of MoyR in the *M. tuberculosis* genome, a 22-bp region; GACGTATTAAAACGTCTTATAC including the consensus sequence of MoyR was searched using FUZZNUC program (Rice, Longden, and Bleasby 2000) and FIMO program (Grant, Bailey, and Noble 2011) using the whole genome sequence of *M. tuberculosis* as the target sequence. There were no matches found up to the cut off *p*-value 1E-6 where a single match was found at *p*-value 1E-5, this signifies that there is no probable any other binding motifs in the *M*. *tuberculosis* genome other than the MoyR consensus motif in the *moyO* region, indicating the specific role of MoyR in the expression of the Rv0789c-Rv0790-Rv0791c-*moyR* operon and Rv0793 gene cluster.

### Binding of MoyR to the *moyO* intergenic DNA

MoyR expected to control the expression of divergently oriented Rv0789c-Rv0790-Rv0791c-*moyR* operon and Rv0793 by binding the intergenic region (Figure 2A). To demonstrate this 119-bp intergenic region denoted as *moyO* was incubated with MoyR protein and the complex formation was assessed by EMSA. MoyR bound to *moyO* forming three discernible stable complexes, C_1,_ C_2,_ and C_3_ (Figure 3A). At the point of half-maximal conversion of the free DNA, all the three complexes appeared and at sufficiently high MoyR concentrations complex C_1_ disappeared, and all DNA was bound as complex C_2_ and C_3_ (lane 15-18, Figure 3A). The difference in migration between complex C_1_, C_2_, and C_3_ indicates that there are more than two MoyR dimers bound to *moyO* in complex C_2_ and C_3_. Percentage complex formation as a function of [MoyR] was fitted to the Hill equation in which the calculated apparent macroscopic dissociation constant (K_d,app_) of the binding was 3.47 nM (Figure3B). This K_d_ likely represents an upper limit as conditions for formation of complexes are nearly stoichiometric (K_d_ ⁓ [DNA]). Binding reaction between MoyR and *moyO* is moderately cooperative which was indicated by Hill coefficient n = 1.2 ± 0.4. Stoichiometry of the binding was calculated *via* the graph of percentage of complex formation versus [MoyR]/ [*moyO*] which showed that three MoyR dimers bind to the intergenic *moyO* region (Figure 3C) and this is consistent with the formation of three complexes in a positive cooperativity. To determine the binding specificity of the reaction, the *moyO* binding region was incubated with increasing concentration of non-specific linearized pET28a+ plasmid keeping the MoyR concentration constant. Complex formation between MoyR and *moyO* was not affected by addition of non-specific pET28a as shown by complete retention of the complexes even with >400-fold binding site equivalents of non-specific pET28a (Figure 3D).

**Figure 3:**
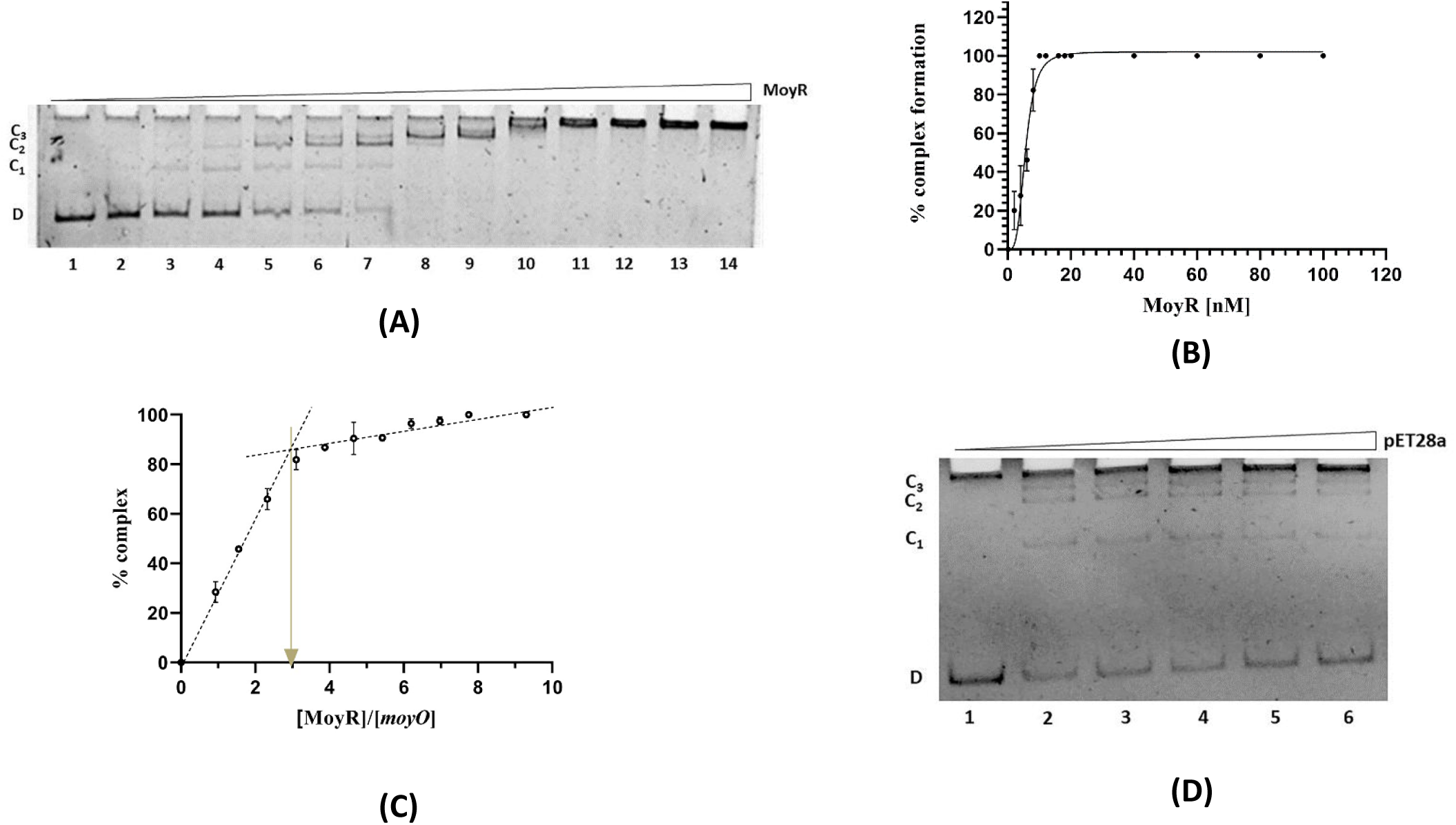
Binding of MoyR to 119 bp intergenic region (*moyO*) of *moyR* and Rv0793 (A) EMSA gel showing binding of MoyR to *moyO* upon increasing concentrations of MoyR (0-100 nM). The stable complexes (C_1_, C_2_, C_3_) and free DNA (D) identified at the left. (B) Percent complex formation of *moyO* region plotted as a function of MoyR concentration. Error bars represent standard deviation (SD) of three independent replicates. (C) Plot of percent complex formation versus [MoyR]/ [*moyO*]. Calculated stoichiometry indicated by the thick arrow points towards the X axis which gives a value of 3 indicating binding of three MoyR dimers to the intergenic region. (D) MoyR (12 nM) binding to *moyO* (3.2 nM) is challenged with increasing concentrations (2-10 nM) of non-specific pET28a plasmid.

**Figure 4:**
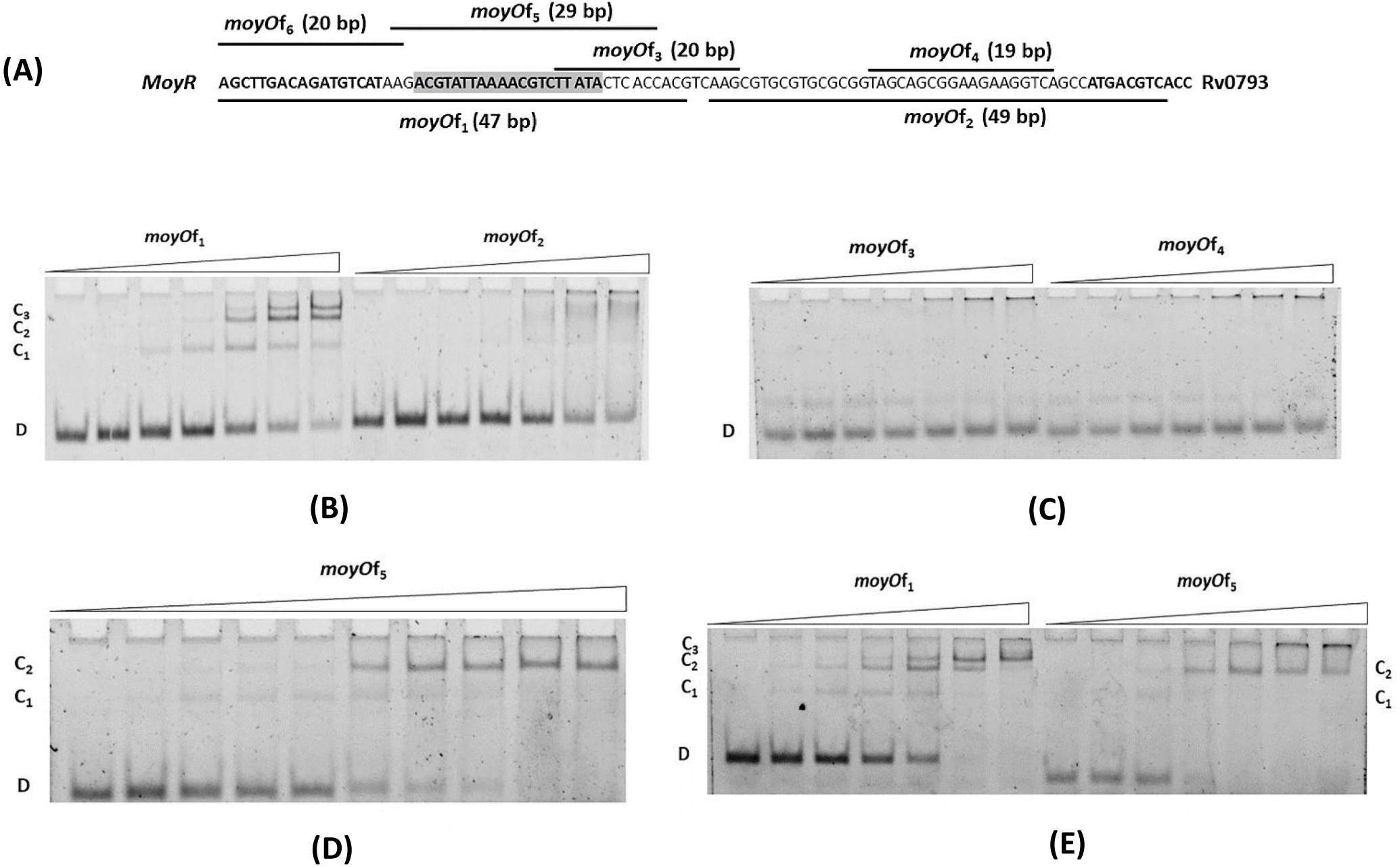
Binding of short oligonucleotides with MoyR. (A) Summary of DNA fragments used in gel mobility shift assays. (B) Electrophoretic mobility shift assay in the presence of *moyO*f_1_ and *moyO*f_2_ with increasing MoyR concentrations ranging from 0 to 100 nM. Binding of MoyR with *moyO*f_1_ formed three stable complexes which is consistent with binding of MoyR with the 119 bp intergenic region. (C) Electrophoretic mobility shift assay in the presence of *moyO*f_3_ and *moyO*f_4_ with increasing MoyR concentrations ranging from 0 to 100 nM. (D) MoyR binding with the identified 19-bp operator sequence. The length of the oligonucleotide (*moyO*f_5_) is 29 bp with extra adjacent base pairs. (E) Qualitative comparison of the migration of MoyR oligomer bound *moyO*f_1_ (47 bp) and *moyO*f_5_ (29 bp). Binding of MoyR with *moyO*f_1_ form three complexes indicating the requirement of a 47 bp region to the formation of the C_3_ complex whereas formation of C_1_ and C_2_ with *moyO*f_5_ specifies that 29 bp region supports only the binding of dimeric and tetrameric MoyR to the DNA.

In order to discern the palindromic binding region more precisely, few shorter DNA oligonucleotides were synthesized and EMSAs were conducted with increasing concentrations MoyR. The DNA fragments were designed in accordance with the *in silico* prediction of *moyO* intergenic region in which *moyO*f_1_ DNA fragment including the predicted MoyR binding motif (Figure 4A). The results showed that MoyR binds to the *moyO*f_1_ fragment forming the complexes C_1_, C_2_, and C_3_ (Figure 4B) with an apparent dissociation constant of 16.6 nM (K_d_ ⁓ [DNA]).

In the presence of *moyO*f_2_ fragment, it showed that MoyR tends to form a very unstable complex at which signifies a nonspecific binding of MoyR with DNA, at high concentrations of protein (Figure 4B). There was no binding observed with the other two shorter DNA fragments, *moyO*f_3_ and *moyO*f_4_ (Figure 4C). As mentioned above, specificity of *moyO*f_1_-MoyR binding was analyzed using linear pET28a+ plasmid and it showed that *moyO*f_1_-MoyR complexes were perpetuated with > 1000-fold excess binding site equivalents of non-specific DNA suggesting that the *moyO*f_1_-MoyR association is specific (data not shown).

The *moyO*f_1_ fragment was further divided to two short fragments, *moyO*f_5_ (29bp) and *moyO*f_6_ (20bp) to assess the binding of MoyR where the *moyO*f_5_ included the predicted 19-bp operator sequence. Results showed that MoyR binds only with *moyO*f_5_ forming only two complexes C_1_ and C_2_ wherein intensity of the bands show that, the formation of C_2_ complex was favoured over formation of C_1_ complex indicating binding of a higher MoyR oligomer than of a dimer (Figure 4D). Migration of the complexes were qualitatively compared upon binding of MoyR with *moyO*f_1_ and *moyO*f_5_ oligonucleotides. Results revealed that migration was similar in C_1_ and C_2_ complexes with both the oligonucleotides indicating binding of similar MoyR oligomeric states to *moyO*f_1_ (47 bp) and *moyO*f_5_ (29 bp) (Figure 4E). The complexes C_1_ and C_2_ correspond to binding of a MoyR dimer and a tetramer respectively. In contrast, C_3_ complex formed with *moyO*f_1_ due to binding of another MoyR dimer to the adjacent nonspecific base pairs other than the palindromic site. Increasing protein concentration with *moyO*f_5_ fragment did not yield any complexes above than the C_2_ complex indicating insufficient length of *moyO*f_5_ for binding of three MoyR dimers (Figure 4E).

### Salt Dependent DNA Binding

As per the results of gel filtration, it clearly showed that MoyR tends to form oligomers depending on the salt concentration in the buffer. Considering the DNA binding region (119 bp), only one possible MoyR binding motif was identified (Figure 4A) indicating the formation of three MoyR-*moyO* complexes due to binding of different MoyR oligomers. To further demonstrate the salt dependency for MoyR-*moyO* binding, an EMSA was carried out with increasing either monovalent or divalent salt concentrations up to 500 mM while maintaining the protein concentration at a constant according to half maximal saturation value. Surprisingly, MoyR-*moyO* complexes appeared to be stable with monovalent salt concentrations much higher than the *M*. *tuberculosis* physiological concentrations (Figure 5A). Increasing monovalent salt concentrations beyond 150 mM has precluded the formation of complex C_1_ and C_3_ whereas complex C_2_ remained stable even at higher concentrations (lane 7-11, Figure 5A). The graph of monovalent salt concentration versus *moyO* fraction bound (curve A, Figure 5C) stipulates that there is no MoyR-*moyO* binding at zero concentration of monovalent cations. Fraction bound less than 1 indicates the poor binding of MoyR to *moyO* in the concentration between 50 to 100 mM whereas fraction bound ⁓ 1 indicates there is no shift of the half maximal saturation point with increasing concentrations of monovalent salts beyond 100 mM. This confirms that there is no dissociation of the complexes but all the protein molecules in the solution were bound to the *moyO* in a higher oligomer state (lane 9-10, Figure 5A). The effect of divalent salts for the binding of MoyR is determined as explained previously for monovalent salts. Increasing concentrations of MgCl_2_ or CaCl_2_ was added to the equilibration buffer without the monovalent salts. Results of the EMSA revealed that, there is no detectable complex formation even at higher concentrations of divalent salts (lane 1-6, Figure 5B).

**Figure 5:**
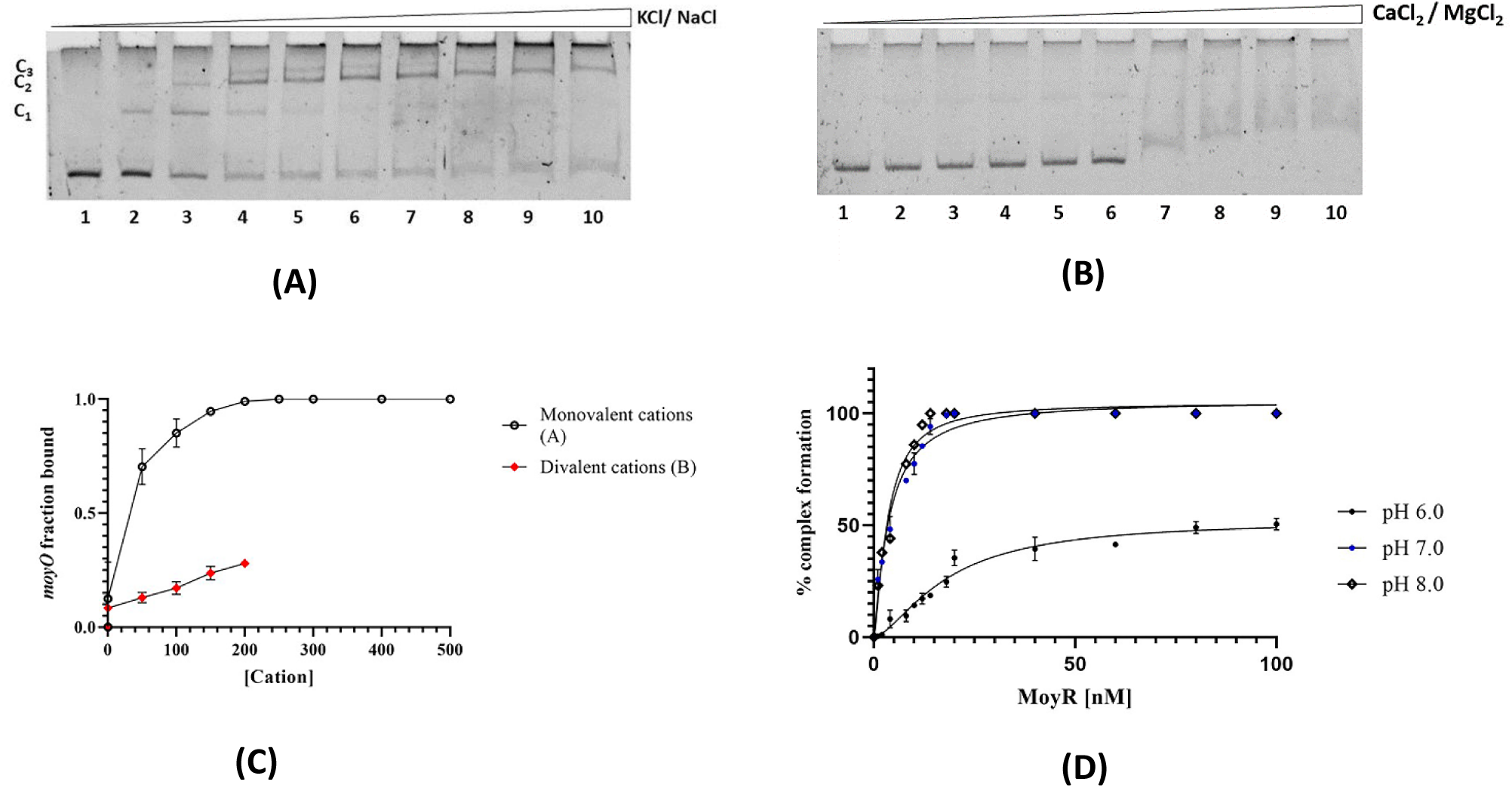
Salt dependent DNA binding of MoyR. (A) MoyR-*moyO* complex formation with increasing KCl/ NaCl (0-500 mM). Lane 3-5 shows 3 distinct stable complexes related to binding of three different MoyR oligomer types, lane 6-8 binding of two oligomer types and lane 9-10 binding of one type of oligomer to *moyO*. (B) MoyR-*moyO* complex formation with increasing CaCl_2_/ MgCl_2_. Lane 2-6 shows a faded band corresponding to complex C_1_, lane 7-10 shows that DNA was degenerated due to high salt. (C) Graph of cation concentration versus *moyO* fraction bound. Last four data points were not included in the curve, wherein the DNA has degenerated due to high salt. (D) Fractional complex formation at pH 8, pH 7 and pH 6 at 100 mM KCl concentration.

According to the graph of divalent cation concentration versus *moyO* fraction bound implies a weak association of MoyR with *moyO*. The fraction bound value remained constant in the range of 0.1 to 0.3, well below the half maximal saturation value (curve B, Figure 5C) indicating substandard formation of MoyR*-moyO* complexes. This suggests that formation of MoyR*-moyO* complexes primarily governed by the monovalent cation concentration in the binding buffer and high concentrations of monovalent salt leads to tight binding of higher oligomers to DNA.

With respect to the three visible stable *MoyR-moyO* complexes (C_1_, C_2_, C_3_) yielded in EMSA macroscopic dissociation constants for complex formation (K_d1_, K_d2_, K_d3_) were assessed at three KCl concentrations, 50 mM, 100 mM and 300 mM. The calculated apparent macroscopic dissociation constants are given by Table 1. These values indicate that there is no significant difference between K_d1_, K_d2_ and K_d3_ at any considered particular KCl. Taken into consideration, binding of MoyR to the *moyO*f_1_ fragment upon increasing KCl triggered the binding of higher oligomers to *moyO*f_1_ as same with the *moyO* region. It clearly denotes that affinity of MoyR to the 119 bp *moyO* intergenic region and 47 bp *moyO*f_1_ fragment remained the same, despite of the fragment length. Further, binding buffers with three different pHs, pH 7.0, pH 6.0 and pH 5.5 were used to determine the influence on pH for MoyR-DNA binding, maintaining the KCl level at a constant, 100 mM concentration. The yielded apparent dissociation constant for the complex formation at pH 7 was 3.7 ± 0.5 nM. Whereas in the binding buffer of pH 6, binding of MoyR to *moyO* was significantly attenuated yielding a K_d,app_ value of 18.4 nM (Figure 5D). In the pH of 5.5, binding of MoyR to *moyO* was hindered wherein a K_d_ value cannot be determined as the unbound DNA remained consistent with increasing protein (data not shown). Considering the data, it reveals that KCl/ NaCl driven MoyR binding to *MoyO* is justifiable only in the pH range of pH 6.0 to pH 8.0 and the binding affinity was attenuated by the pH as well.

**Table 1:** Calculated apparent macroscopic dissociation constants for different complex formation at different KCl monovalent cation concentrations. The value K_d1_ indicates the dissociation constant considering ‘C_1_, C_2_, C_3_’ as ‘one complex’ (complex E_1_) whereas K_d2_ indicates the dissociation constant considering ‘C_2_ and C_3_’ as ‘one complex’ (complex E_2_). Then the K_d3_ was determined considering only C_3_ as the complex (complex E_3_).

| *KCl concentration (mM)* | *K<sub>d1</sub>(complex E<sub>1</sub>)* | *K<sub>d2</sub>(complex E<sub>2</sub>)* | *K<sub>d3</sub>(complex E<sub>3</sub>)* |
| --- | --- | --- | --- |
| 50 | 6.4 nM | 7.6 nM | 10.7 nM |
| 100 | 3.5 nM | 3.5 nM | 3.4 nM |
| 300 | 2.7 nM | 2.9 nM | 3.8 nM |

### pH Dependent Protonation-deprotonation of MoyR

Transcriptional regulators are sensitive to the pH of the environment, and the pH dependence can be attributed to the protonation-deprotonation of the functional groups of amino acids thereby affecting the dimerization and conformational changes in the proteins. Taken into consideration, pKa values of the titratable residues of MoyR were identified using using H++ server and PlayMolecule server (Gordon *et al*., 2005; Martinez-rosell *et al*., 2017). Out of 269 amino acids in the monomeric MoyR, there were 57 titratable residues naming cysteine, lysine, arginine, aspartic acid, histidine, and tyrosine. Results revealed protonation of amino acids needed for the protein-DNA binding, indicating a higher number of protonated residues than deprotonated residues at pH 8.0 and pH 7.0 where 31 were protonated. One dubious residue Cys95 was identified at pH 8, with a pKa of 8.29. It showed that in addition to the 31 protonated residues, another three residues were protonated at pH 5.0, Cys95, Glu92 and Glu243 with three dubious residues Glu90 (pKa = 5.52), His195 (pKa = 5.05) and His239 (pKa = 5.37). There were no significant pKa value changes in the conserved DNA binding residues upon decreasing pH from pH 8.0 to 5.0. This indicates that DNA binding residues of MoyR do not undergo any protonation-deprotonation changes within the pH range 8 to 5. Nevertheless, conserved His195 in the ligand binding domain was subjected to deprotonation at pH 5.0 from protonated state at pH 8.0. This implies that His195 residue plays a major role in pH sensitivity of MoyR in relation to oligomer stability of the protein.

To estimate the oligomer formation and aggregation of MoyR, particle sizes of MoyR were analyzed using dynamic light scattering (DLS) at different pHs varied from pH 7.5 to pH 5.5. The results of DLS analysis revealed a relationship between that particle size of MoyR at different pHs. Comparison of hydrodynamic radius and molecular weight in different buffers were shown in Figure 6A. According to the obtained measurements MoyR hydrodynamic radius was minimum in the pH 5.5 buffer with a monodisperse size distribution whereas different hydrodynamic radii were obtained at pH 6.0 and above with a polydisperse size distribution. Particle size is more similar in the pH range 7.5 to 6.0 suggesting similar oligomer states within this pH range. There was no aggregation observed. High light scattering indicated aggregation of MoyR in the presence of monovalent cations in the pH 5.5 buffer (Figure 6B). This indicated that MoyR tends to form aggregates in the presence of monovalent cations in low pHs which can lead to inhibition of its DNA binding due to aggregation in low pH values. This was corroborated with the K_d_ values obtained under different pHs, where MoyR-DNA binding was hindered at pH 5.5. Therefore, it is evident that, monovalent cation driven oligomerization and DNA binding can be justifiable only in the physiological pH range.

**Figure 6:**
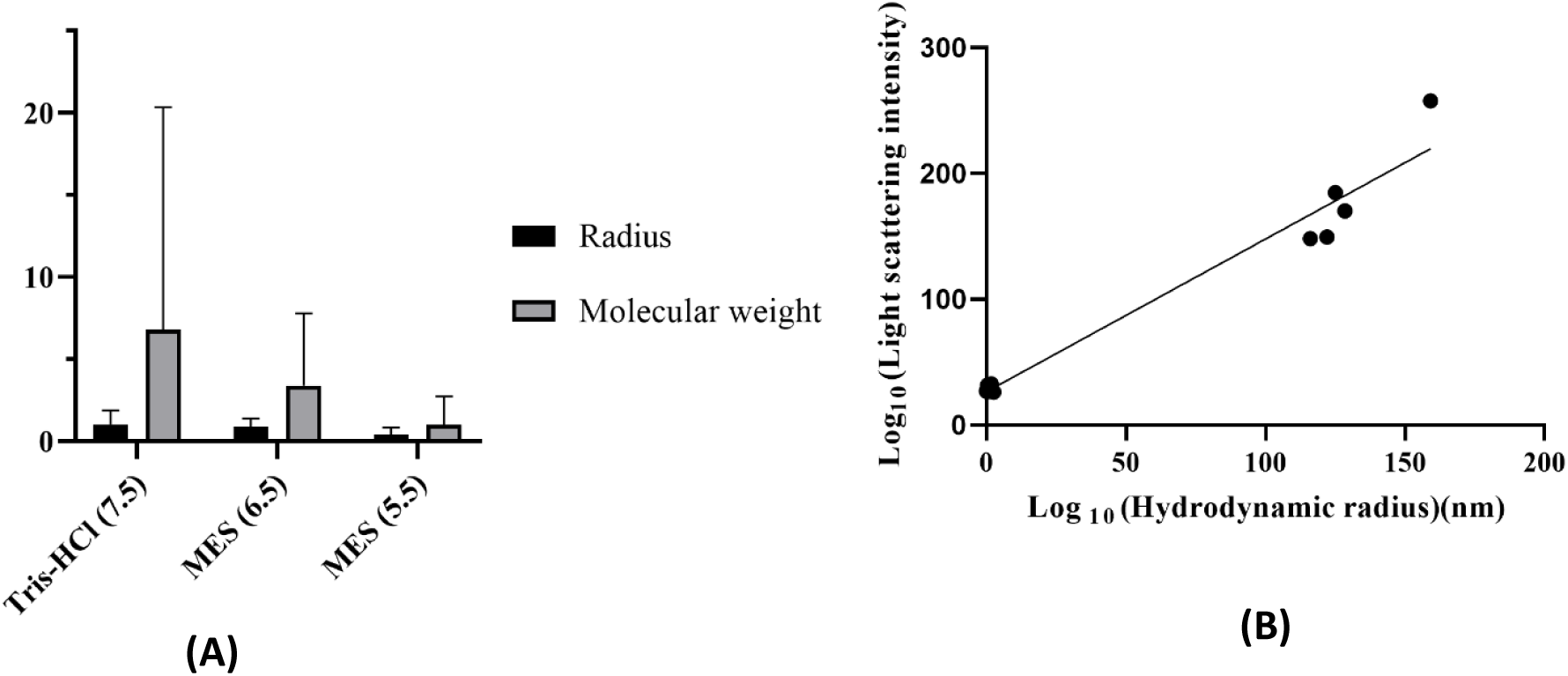
Estimating the particle sizes of MoyR in different pH buffers. (A) Graph of particle radius and molecular weight of MoyR oligomers in pH 7.5, pH 6.5 and pH 5.5. The lowest particle size and the molecular weight observed at pH 5.5 indicating the presence of monomeric MoyR whereas the distribution of oligomers varying from lowest to higher molecular weights were observed at pH 6.5 and above. (B) Graph of light scattering intensity versus hydrodynamic radius (nm) to differentiate oligomers and aggregates at pH 5.5. Two clustered data points showing the lower radius corresponding to monomers, dimers, tetramers etc without monovalent cations, whereas the aggregates showed a radius 100 times greater than of the oligomeric MoyR in the presence of monovalent cations.

**Figure 7:**
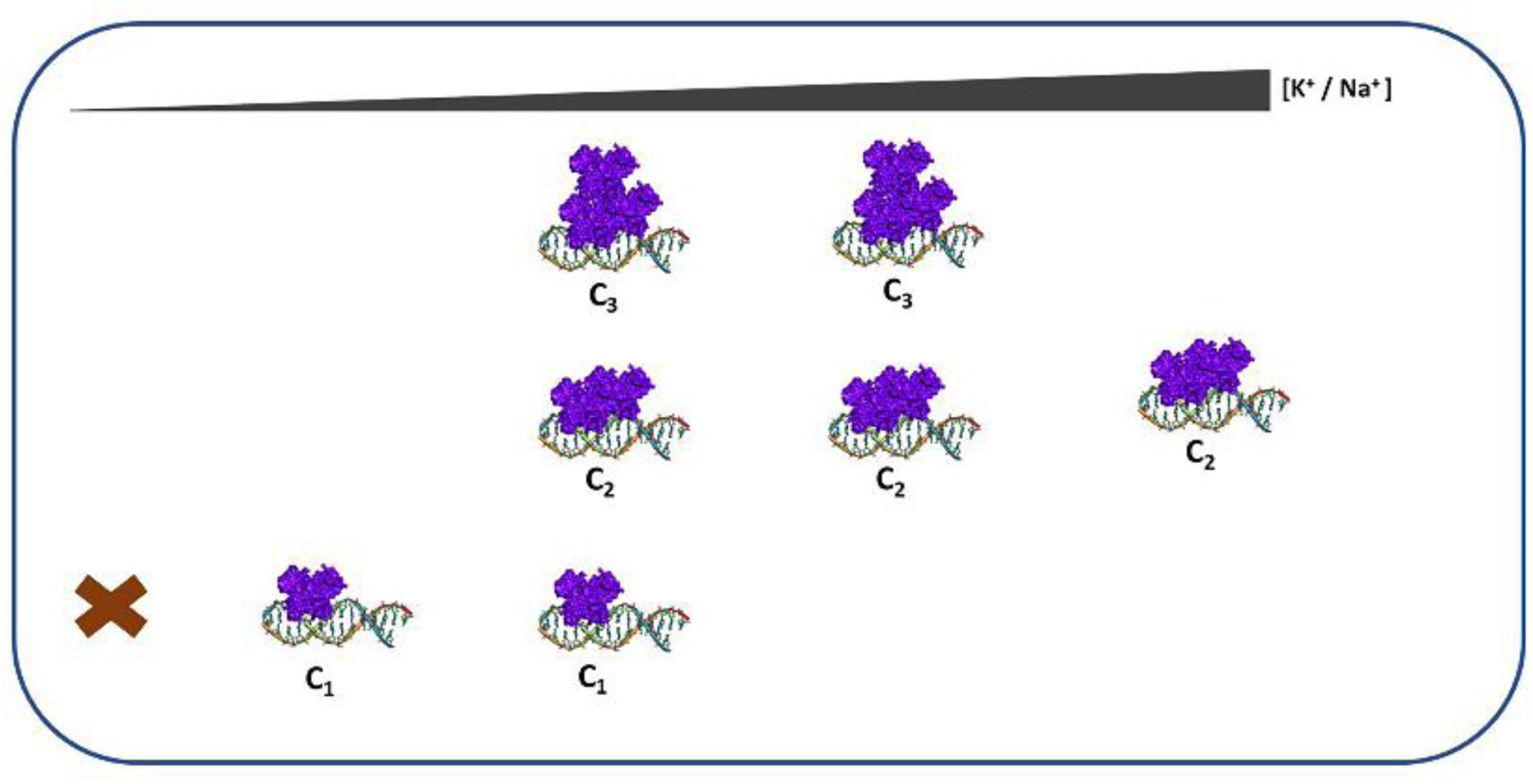
Schematic representation of the proposed model for MoyR interaction with the operator sequence. MoyR oligomer stability upon increasing K^+^ / Na^+^ concentrations. In moderate K^+^ / Na^+^ concentration MoyR dimer (C_1_), tetramer (C_2_) and hexamer (C_3_) independently bind to the operator sequence. Upon high concentrations of K^+^ / Na^+^, MoyR dimer and hexamer transition to tetramer take place wherein tetramer bound complex remained stable with a high affinity.

### Probing for a natural ligand

Further, a variety of natural ligands act as effectors for HutC regulators, including histidine, sugar-phosphates, long-chain fatty acids etc. With accordance to the adjacent hypothetical monoxygense and oxidoreductase encoding genes in the *moyR* gene locus (Figure 3A) various possible pathways can attribute with MoyR, as monooxygenases and oxidoreductases are involved in many cellular processes. According to the instances of gene expression of the *moyR* gene cluster in the presence of different fatty acids, pyrazinamide, hypoxia and supported data by literature, a hypothesis can be made that, MoyR can play a regulatory role in fatty acid synthesis pathway or polyketide antibiotic synthesis pathway. A molecular simulation approach was used to deduce the possible ligands that can bind with MoyR, using intermediate compounds in cellular pathways such as, fatty acid synthesis pathways and polyketides biosynthesis pathways, krebs cycle, glyoxylate cycle, chorismate synthesis and cholesterol synthesis due to the noteworthy contribution of monooxygenases and oxidoreductases in these pathways. Notably, intermediate compounds in type II polyketide biosynthesis pathway showed a prominent binding with MoyR than the compounds of fatty acid synthesis pathway and other pathways. Many of these intermediates act as starting compounds for synthesis of various secondary compounds that are therapeutically important such as actinorhodin, tetracycline, and mithramycin. Surprisingly, the majority of the highest affinity compounds with lower binding energies were the intermediate compounds involved in tetracycline biosynthesis (Table 2). Molecular docking of of Rv0791 and Rv0793 revealed a similar pattern in binding with high affinity to the intermediates of type II polyketide synthesis including landomycin, granaticin, tetracycline and chlorotetracycline.

**Table 2:**
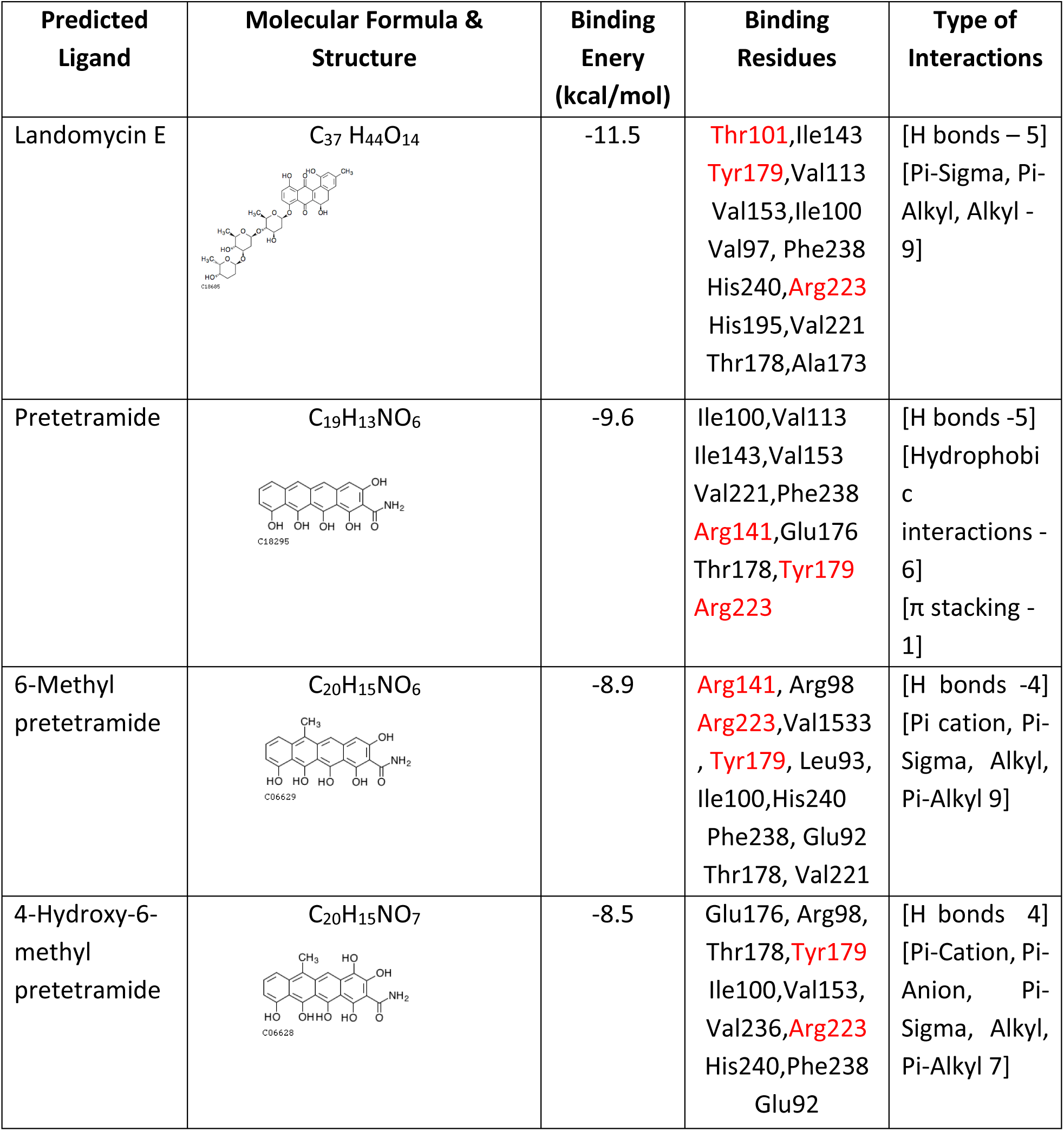
Predicted high affinity natural ligands for MoyR. Conserved important binding residues in the binding pocket are highlighted in red.

| Predicted Ligand | Molecular Formula & Structure | Binding Enery (kcal/mol) | Binding Residues | Type of Interactions |
| --- | --- | --- | --- | --- |
| Landomycin E | **C<sub>37</sub> H<sub>44</sub> O<sub>14</sub>**<br>C18685 | -11.5 | Thr101,Ile143<br>Tyr179,Val113<br>Val153,Ile100<br>Val97, Phe238<br>His240,Arg223<br>His195,Val221<br>Thr178,Ala173 | [H bonds – 5]<br>[Pi-Sigma, Pi-Alkyl, Alkyl - 9] |
| Pretetramide | **C<sub>19</sub>H<sub>13</sub>NO<sub>6</sub>**<br>C18295 | -9.6 | Ile100,Val113<br>Ile143,Val153<br>Val221,Phe238<br>Arg141,Glu176<br>Thr178,Tyr179<br>Arg223 | [H bonds -5]<br>[Hydrophobic interactions - 6]<br>[π stacking - 1] |
| 6-Methyl pretetramide | **C<sub>20</sub>H<sub>15</sub>NO<sub>6</sub>**<br>C06629 | -8.9 | Arg141, Arg98<br>Arg223,Val1533<br>, Tyr179, Leu93,<br>Ile100,His240<br>Phe238, Glu92<br>Thr178, Val221 | [H bonds -4]<br>[Pi cation, Pi-Sigma, Alkyl, Pi-Alkyl 9] |
| 4-Hydroxy-6-methyl pretetramide | **C<sub>20</sub>H<sub>15</sub>NO<sub>7</sub>**<br>C06628 | -8.5 | Glu176, Arg98,<br>Thr178,Tyr179<br>Ile100,Val153,<br>Val236,Arg223<br>His240,Phe238<br>Glu92 | [H bonds 4]<br>[Pi-Cation, Pi-Anion, Pi-Sigma, Alkyl, Pi-Alkyl 7] |

None of the sugar moieties, intermediates of fatty acid biosynthesis, krebs cycle intermediates or chorismate synthesis showed a considerable affinity for any of these proteins, MoyR, Rv0791 and Rv0793. This implies that MoyR protein could be playing a role in the type II polyketide biosynthesis, specially, including tetracycline biosynthesis which need further investigations.

## Discussion

Transcriptional regulators play a crucial role wherein they control the cellular mechanisms by regulation of gene expression accordingly in the presence or absence of small molecules such as metabolites, ions and drug molecules. Bacteria possess many transcriptional regulators and the GntR family of wHTH-type transcriptional regulators are an important class of proteins involved in diverse roles in the bacteria including primary metabolism, virulence, development, antibiotic resistance, antibiotic production etc (Hoskisson and Rigali 2009). In this context, transcriptional regulators including the GntR family, have drawn significant attention as drug targets for antibiotic-resistant pathogenic bacteria such as *M*. *tuberculosis*. Hence, this study was focused on characterizing one of the GntR regulators MoyR in the *M*. *tuberculosis* genome. The genomic locus of MoyR consists of many hypothetical proteins, data from our previous study together with previous published literature suggested that adjacent genes of MoyR encode proteins are homologous to different monooxygenases and it is highly likely that MoyR regulates the expression of a gene cluster that encodes for different monooxygenases (Abeywickrama and Perera 2021; Sciara et al. 2003; Lemieux et al. 2005; Glasser et al. 2017). The global regulator DasR from *S*. *coelicolor* was identified as the structural homologue of MoyR and shared a considerable sequence similarity with both DasR and NagR from *Bacillus subtilis* which are well known HutC regulators. The operator sequence *dre-*sites of DasR have been characterized in great detail wherein NagR also recognizes the *dre*-site for binding (Colson et al. 2006; Fillenberg et al. 2015). Multiple *dre*-sites have been identified in the genome of *S*. *coelicolor* for DasR and *B*. *subtilis* for NagR signifying the global regulatory role of the proteins. However, the analysis of the *moyR*-Rv0793 intergenic region of MoyR did not reveal any *dre*-sites. Instead, a different HutC motif was identified in the intergenic region, that shares a high similarity with the binding motif of fatty acyl responsive regulator FarR from *E*. *coli*. In the present study we identified only one MoyR binding site in the upstream region of *moyR* and Rv0793. However, there were no other MoyR binding motifs identified in the genome of *M*. *tuberculosis,* and this endorses that MoyR regulates a group of monooxygenases related to a specific cellular pathway of the bacteria. Considering the MoyR binding with its intergenic region, the obtained dissociation constant values for MoyR-*moyO* complex formation were in the nanomolar range, indicating a considerable high affinity of MoyR to the binding site and this implies that MoyR is a functional transcriptional regulator in *M*. *tuberculosis*. Notably, three distinct stable MoyR-*moyO* complexes were formed indicating binding of multiple MoyR dimers to the binding site with a high affinity. The formation of the three distinct complexes were observed with the short oligonucleotide (*moyO*_f1_) that consists of the specific binding site, hence the nonspecific binding of dimers into the intergenic region can be clearly excluded. Results of gel filtration indicated that MoyR is predominantly a dimer in solution at physiologically relevant monovalent cation concentrations, but at 500 mM KCl/ NaCl concentration MoyR tends to form oligomers such as tetramers and hexamers. This was further corroborated by the analysis of the DSC thermogram of MoyR that it was well fitted to the non-two-state model. This implies that folding of MoyR in the solution occurs via stable intermediates or there can be different oligomeric states of MoyR. The obtained enthalpy Δ*H*_cal_ was 76.8 ± 16.3 kcal/mol and Δ*H*_vant_ was 127.9 ± 33.6 kcal/mol in which Δ*H*_cal_/Δ*H*_vant_ is roughly 0.5 suggesting that MoyR has a dimeric assembly in solution. Surprisingly, two out of the three MoyR-*moyO* complexes were stable upon increasing K^+^/ Na^+^ concentrations up to 300 mM, whereas one complex was stable even up to 500 mM K^+^/ Na^+^ concentration. Strikingly, the MoyR-DNA complex stability remained up to 1 M K^+^ (data not shown) which is unusual for a typical protein-DNA binding. Moreover, size exclusion chromatography, glutaraldehyde crosslinking and *in silico* analysis provide strong evidence that MoyR has a higher tendency to form oligomers which are greatly induced by the presence of K^+^ and Na^+^ in the medium. Hence, we suggest that the three MoyR-*moyO* complexes represent the binding of a MoyR dimer, a tetramer, and a hexamer with the operator sequence. Taken together, herein we propose a model for MoyR in regulating *moyO* promoter activities which involve complex oligomerization in response to the presence of K^+^ and Na^+^ in the binding buffer in the physiologically relevant pH. In very low concentrations of K^+^/ Na^+^ (0-50 mM), the MoyR binding to DNA is restrained, whereas in the moderate K^+^/ Na^+^ (70-150 mM) concentrations MoyR binds to DNA in all three forms: as a dimer, tetramer and a hexamer yielding different molar weight complexes. In contrast, upon high K^+^/ Na^+^ (200-500 mM) concentrations only the tetramer remained bound to the DNA whereas the dimer and hexamer dissociate from the cognate DNA to form the tetramer (Figure 6). This is further confirmed by the amount of free DNA available in lanes 4-10 in Figure 5A, and the half maximal saturation point remained constant while the bound MoyR switched the oligomeric state from dimer/ hexamer to tetramer.

The localization of a transcription factor to a promoter depends on the dimerization of the transcription factor because DNA binding domains of most factors are palindromic, recognizing half-sites of the operator sequence. However, for some transcription factors dimerization alone cannot stabilize the binding do DNA. If the dimer has insufficient affinity to be biologically effective, oligomerization of identical monomers can supplement the stability of protein-DNA complexes. Higher oligomeric states of regulators have been predominantly observed in the transcriptional regulator family LysR, which has been extensively studied (Sainsbury et al. 2009; Gong, Xiong, and Maser 2012). Unlike in LysR regulators, although the higher oligomers of GntR regulators can be observed in solution, so far, all the identified regulators function as dimers. Remarkably, a very recent study by Vigouroux *et al* made a landmark in which they proposed the first tetrameric transcription factor in the GntR/ VanR family. The regulator, Atu1419 from the plant pathogen *Agrobacterium fabrum* was shown to be present in the tetrameric form in solution and it binds the DNA as a tetramer which was confirmed by the DNA-bound crystal structure. It has been reported that Atu1419 is a tetramer in its apo form, DNA-bound and effector bound form suggesting its quaternary structure is the biologically active form (Vigouroux et al. 2021).

The surrounding counterions play an important role in the stability of protein and DNA, wherein the counterions accumulate in the vicinity of anionic nucleic acid to neutralize the phosphate charges which would hinder the protein-DNA association by high concentrations of monovalent cations. Commonly in mesophilic bacteria, the protein-DNA complexes are stable in moderate salt concentrations whereas concentrations above 300 mM can dissociate the protein-DNA complexes and this has been extensively studied (Kwon et al. 2001; Šebest et al. 2015; Hart et al. 1999). Halophilic bacteria are an exception, wherein these organisms maintain intracellular salt concentrations close to saturation to survive in high-salt environments. The significant feature of these bacteria is that their adaptation of cellular processes to function under high intracellular salt concentrations nevertheless little is known regarding halophilic adaptation to DNA processing machinery. A notable wide range of salt tolerance from 1% to 8% was reported for different *Mycobacterium* spp in which the bacteria grouped as ‘salt-sensitive’, ‘salt-intermediate’ and ‘salt-tolerance’. It has further shown an inverse correlation between the salt tolerance and the host adaptation in which the environmental *Mycobacterium* spp are more salt tolerant than the pathogenic *Mycobacterium* spp (Asmar et al. 2016). However, *M*. *tuberculosis* encounters electrolyte and osmotic stress within the host which is significantly higher than the culture media wherein concentration of NaCl varies within the respiratory route from 50 mM in airway surface liquid to 250 mM in macrophages(Marino et al. 2016). Initially, it was shown that mycobacteria experience a K^+^ deficiency in the phagosome, but in a recent study, it has been elicited that intraphagosomal K^+^ increases during macrophage phagosome maturation that K^+^ is not limiting during macrophage infection. This clearly indicates that *M*. *tuberculosis* is exposed to various extracellular Na^+^ and K^+^ concentrations during infection that is significantly different to that of *in vitro* culture media and the intracellular Na^+^ and K^+^ can vary during infection (Macgilvary, Kevorkian, and Id 2019; Salina et al. 2014). In this context, it we can propose that intracellular Na^+^ and K^+^ levels can directly remodel the oligomerization of MoyR protein and thereby regulate the gene expression. Furthermore, MoyR tetramer association with the DNA is compatible under intracellular conditions causing tight repression of the gene cluster. The results of MoyR-*moyO* binding at various pH levels indicated that the tight association of MoyR and the DNA occurred in the pH range 6 to 8 wherein mycobacteria maintain intracellular pH between 6.1 to 7.2 (Rao et al. 2001). High thermal stability and high affinity to DNA upon high monovalent salts manifest that MoyR repressor activity can be remained even under unfavourable cellular environments. Moreover, it is paramount to investigate the MoyR oligomer-bound DNA stability upon high concentrations of salt and its physiological relevance to the bacterium.

## Conclusion

There are evidences that *moyR* (Rv0792c) is highly expressed in hypoxic, oxidative stress conditions and when the bacterium persists within the phagosomal environment (Schnappinger et al. 2003; Arun et al. 2018; Rustad et al. 2008; Sun et al. 2018). Herein we have identified MoyR high thermal stability and high affinity to DNA upon high monovalent cations, which is unusual for a mesophilic bacterium and this manifest that MoyR repressor activity can be remained even under unfavorable cellular environments. Hence, this denotes that MoyR can play an important role during infection and persistence where the bacterium encounters extreme stress conditions. The early progenitor of *Mycobacterium* is an environmental bacterium, and this evokes that the *M*. *tuberculosis* genome still consists of cellular pathways responsible for surviving in environments such as soil and water other than animals. Furthermore, it is considered that the *Mycobacterium* genome is more conserved among the species. This indicates that there could be inactive antibiotic synthesis pathways or similar pathways still in the mycobacterial genome and this could be the genesis of antibiotic resistance in the bacterium. Recent molecular biology studies of *Streptomyces* and *Mycobacterium* have revealed prominent similarities in the developmental and morphological characteristics of the two bacteria. One simple example is the striking similarities of the crystal structure of ActVA-Orf6 from *Streptomyces* with the Rv0793 of *M*. *tuberculosis* wherein ActVA-Orf6 participates in a polyketide antibiotic synthesis pathway.

Thus, it can predict that there could be similar polyketide synthesis pathways shared by both the bacteria. This brings forth the importance of MoyR as a drug target in *M*. *tuberculosis* which could be of great interest if *M*. *tuberculosis* consists of any inactive antibiotic synthesis pathways wherein this could lead to a novel paradigm to the antibiotic tolerance and resistance mechanisms and of the bacterium.

## Acknowledgements

This work was funded by a university grant (grant no-AP/3/2/2014/RG/04), University of Colombo, Sri Lanka. This work was supported by Institute for Protein Research, Osaka University, Japan.

## References

Abeywickrama, Thanusha Dhananji, and Inoka Chinthana Perera. 2021. “In Silico Characterization and Virtual Screening of GntR/HutC Family Transcriptional Regulator MoyR: A Potential Monooxygenase Regulator in Mycobacterium Tuberculosis.” Biology 10 (12): 1241. 10.3390/biology10121241.

Aravind, L., Vivek Anantharaman, Santhanam Balaji, M. Mohan Babu, and Lakshminarayan M. Iyer. 2005. “The Many Faces of the Helix-Turn-Helix Domain: Transcription Regulation and Beyond.” FEMS Microbiology Reviews 29 (2): 231–62. 10.1016/j.femsre.2004.12.008.

Arun, P V Parvati Sai, Sravan Kumar Miryala, Aarti Rana, and Sreenivasulu Kurukuti. 2018. “System-Wide Coordinates of Higher Order Functions in Host-Pathogen Environment upon Mycobacterium Tuberculosis Infection.” Scientific Reports 8: 1–12. 10.1038/s41598-018-22884-8.

Asmar, Shady, Mohamed Sassi, Michael Phelippeau, and Michel Drancourt. 2016. “Inverse Correlation between Salt Tolerance and Host-Adaptation in Mycobacteria.” BMC Research Notes 9 (1): 1–9. 10.1186/s13104-016-2054-y.

Bailey, Timothy L., Mikael Boden, Fabian A. Buske, Martin Frith, Charles E. Grant, Luca Clementi, Jingyuan Ren, Wilfred W. Li, and William S. Noble. 2009. “MEME Suite: Tools for Motif Discovery and Searching.” Nucleic Acids Research 37 (SUPPL. 2): 202–8. 10.1093/nar/gkp335.

Biswas, Rajesh Kumar, Debashis Dutta, Ashutosh Tripathi, Youjun Feng, Monisha Banerjee, and Bhupendra N. Singh. 2013. “Identification and Characterization of Rv0494: A Fatty Acid-Responsive Protein of the GntR/FadR Family from Mycobacterium Tuberculosis.” Microbiology (United Kingdom) 159 (PART 5): 913–23. 10.1099/mic.0.066654-0.

Colson, Séverine, Joachim Stephan, Tina Hertrich, Akihiro Saito, Gilles P. Van Wezel, Fritz Titgemeyer, and Sébastien Rigali. 2006. “Conserved Cis-Acting Elements Upstream of Genes Composing the Chitinolytic System of Streptomycetes Are DasR-Responsive Elements.” Journal of Molecular Microbiology and Biotechnology 12 (1–2): 60–66. 10.1159/000096460.

Crooks, G, G Hon, J Chandonia, and S Brenner. 2004. “NCBI GenBank FTP Site\nWebLogo: A Sequence Logo Generator.” Genome Res 14: 1188–90. 10.1101/gr.849004.1.

Dallakyan, S., & Olson, A. J. (2015). Small-Molecula Library Screening by Docking with PyRx. Methods in Molecular Biology, 243–250.

Fillenberg, Simon B., Florian C. Grau, Gerald Seidel, and Yves A. Muller. 2015. “Structural Insight into Operator Dre-Sites Recognition and Effector Binding in the GntR/HutC Transcription Regulator NagR.” Nucleic Acids Research 43 (2): 1283–96. 10.1093/nar/gku1374.

Franco, Irina Saraiva, Luís Jaime Mota, Cláudio Manuel Soares, and Isabel De Sá-Nogueira. 2006. “Functional Domains of the Bacillus Subtilis Transcription Factor AraR and Identification of Amino Acids Important for Nucleoprotein Complex Assembly and Effector Binding.” Journal of Bacteriology 188 (8): 3024–36. 10.1128/JB.188.8.3024-3036.2006.

Fujita, Y., T. Fujita, Y. Miwa, J. Nihashi, and Y. Aratani. 1986. “Organization of Transcription of the Gluconate Operon, Gnt, of Bacillus Subtilis.” Journal of Biological Chemistry 261 (29): 13744–53. 10.1016/s0021-9258(18)67083-8.

Gagneux, Sebastien. 2018. “Ecology and Evolution of Mycobacterium Tuberculosis.” Nature Reviews Microbiology. 10.1038/nrmicro.2018.8.

Gajiwala, Ketan S., and Stephen K. Burley. 2000. “Winged Helix Proteins.” Current Opinion in Structural Biology 10 (1): 110–16. 10.1016/S0959-440X(99)00057-3.

Glasser, Nathaniel R., Benjamin X. Wang, Julie A. Hoy, and Dianne K. Newman. 2017. “The Pyruvate and α-Ketoglutarate Dehydrogenase Complexes of Pseudomonas Aeruginosa Catalyze Pyocyanin and Phenazine-1-Carboxylic Acid Reduction via the Subunit Dihydrolipoamide Dehydrogenase.” Journal of Biological Chemistry 292 (13): 5593–5607. 10.1074/jbc.M116.772848.

Gong, Wenjie, Guangming Xiong, and Edmund Maser. 2012. “Oligomerization and Negative Autoregulation of the LysR-Type Transcriptional Regulator HsdR from Comamonas Testosteroni.” Journal of Steroid Biochemistry and Molecular Biology 132 (3–5): 203–11. 10.1016/j.jsbmb.2012.05.012.

Grant, Charles E., Timothy L. Bailey, and William Stafford Noble. 2011. “FIMO: Scanning for Occurrences of a given Motif.” Bioinformatics 27 (7): 1017–18. 10.1093/bioinformatics/btr064.

Hart, Darren J, Robert E Speight, Matthew A Cooper, John D Sutherland, and Jonathan M Blackburn. 1999. “The Salt Dependence of DNA Recognition by NF-κ B P50: A Detailed Kinetic Analysis of the Effects on Affinity and Specificity.” Nucleic Acids Research 27 (4): 1063–69.

Hellman, Lance M., and Michael G. Fried. 2007. “Electrophoretic Mobility Shift Assay (EMSA) for Detecting Protein-Nucleic Acid Interactions.” Nature Protocols 2 (8): 1849–61. 10.1038/nprot.2007.249.

Hoskisson, Paul A., and Sébastien Rigali. 2009. Variation in Form and Function, The Helix-Turn-Helix Regulators of the GntR Superfamily. Advances in Applied Microbiology. Vol. 69. 10.1016/S0065-2164(09)69001-8.

Hoskisson, Paul A., Sebastien Rigali, Kay Fowler, Kim C. Findlay, and Mark J. Buttner. 2006. “DevA, a GntR-like Transcriptional Regulator Required for Development in Streptomyces Coelicolor.” Journal of Bacteriology 188 (14): 5014–23. 10.1128/JB.00307-06.

Kwon, Haeyoung, Seyeon Park, Sangkyou Lee, Dug-keun Lee, and Chul-hak Yang. 2001. “Determination of Binding Constant of Transcription Factor AP-1 and DNA Application of Inhibitors.” Eur. J. Biochem 268: 565–72.

Lee, Martin H., Michael Scherer, Sébastien Rigali, and James W. Golden. 2003. “PlmA, a New Member of the GntR Family, Has Plasmid Maintenance Functions in Anabaena Sp. Strain PCC 7120.” Journal of Bacteriology 185 (15): 4315–25. 10.1128/JB.185.15.4315-4325.2003.

Lemieux, M. Joanne, Claire Ference, Maia M. Cherney, Metian Wang, Craig Garen, and Michael N.G. James. 2005. “The Crystal Structure of Rv0793, a Hypothetical Monooxygenase from M. Tuberculosis.” Journal of Structural and Functional Genomics 6 (4): 245–57. 10.1007/s10969-005-9004-6.

Macgilvary, Nathan J, Yuzo L Kevorkian, and Shumin Tan Id. 2019. “Potassium Response and Homeostasis in Mycobacterium Tuberculosis Modulates Environmental Adaptation and Is Important for Host Colonization.” PLoS Pathogens 15 (2): 1–23.

Marino, Leonardo B, Ashoka V R Madduri, Δ T J Ragan, Debbie M Hunt, Lucrezia Bassano, Maximiliano G Gutierrez, D Branch Moody, Fernando R Pavan, and Luiz Pedro S De Carvalho. 2016. “Cell-Envelope Remodeling as a Determinant of Phenotypic Antibacterial Tolerance in Mycobacterium Tuberculosis.” ACS Infectious Diseases 2: 352–60. 10.1021/acsinfecdis.5b00148.

Morris, G. M. (2010). AutoDock Version 4.2 - User Guide. 1–49.

Ó’Fágáin, Ciarán, Philip M. Cummins, and Brendan O’Connor. 2017. “Gel-Filtration Chromatography.” Methods in Molecular Biology 1485: 15–25. 10.1007/978-1-4939-6412-3_2.

Perera, Inoka C., and Anne Grove. 2010. “Urate Is a Ligand for the Transcriptional Regulator PecS.” Journal of Molecular Biology 402 (3): 539–51. 10.1016/j.jmb.2010.07.053.

Perez, Jose maria Mateos, and Javier Pascau. 2013. Image Processing with ImageJ. Edited by chavarris Cristina. PACKT Publishing.

Pettersen, E., Goddard, T., Huang, C., Couch, G., Greenblatt, D., Meng, E., & Ferrin, T. (2004). UCSF Chimera--a visualization system for exploratory research and analysis. J Comput Chem, 25(13), 1605–1612.

Ranganathan, Sridevi, Jonah Cheung, Michael Cassidy, Christopher Ginter, Janice D. Pata, and Kathleen A. McDonough. 2018. “Novel Structural Features Drive DNA Binding Properties of Cmr, a CRP Family Protein in TB Complex Mycobacteria.” Nucleic Acids Research 46 (1): 403–20. 10.1093/nar/gkx1148.

Rao, Min, Trevor L Streur, Frank E Aldwell, and Gregory M Cook. 2001. “Intracellular PH Regulation by Mycobacterium Smegmatis and Mycobacterium Bovis BCG.” Microbiology 147: 1017–24.

Rice, Peter, Lan Longden, and Alan Bleasby. 2000. “EMBOSS: The European Molecular Biology Open Software Suite.” Trends in Genetics 16 (6): 276–77. 10.1016/S0168-9525(00)02024-2.

Rigali, S. 2004. “Extending the Classification of Bacterial Transcription Factors beyond the Helix-Turn-Helix Motif as an Alternative Approach to Discover New Cis/Trans Relationships.” Nucleic Acids Research 32 (11): 3418–26. 10.1093/nar/gkh673.

Rigali, Sébastien, Adeline Derouaux, Fabrizio Giannotta, and Jean Dusart. 2002. “Subdivision of the Helix-Turn-Helix GntR Family of Bacterial Regulators in the FadR, HutC, MocR, and YtrA Subfamilies.” Journal of Biological Chemistry 277 (15): 12507–15. 10.1074/jbc.M110968200.

Rustad, Tige R., Maria I. Harrell, Reiling Liao, and David R. Sherman. 2008. “The Enduring Hypoxic Response of Mycobacterium Tuberculosis.” PLoS ONE 3 (1): 1–8. 10.1371/journal.pone.0001502.

Sainsbury, Sarah, Laura A. Lane, Jingshan Ren, Robert J. Gilbert, Nigel J. Saunders, Carol V. Robinson, David I. Stuart, and Raymond J. Owens. 2009. “The Structure of CrgA from Neisseria Meningitidis Reveals a New Octameric Assembly State for LysR Transcriptional Regulators.” Nucleic Acids Research 37 (14): 4545–58. 10.1093/nar/gkp445.

Salina, Elena G, Simon J Waddell, Nadine Hoffmann, Ida Rosenkrands, Philip D Butcher, and Arseny S Kaprelyants. 2014. “Potassium Availability Triggers Mycobacterium Tuberculosis Transition to, and Resuscitation from, Non-Culturable (Dormant) States.” Open Biology 4.

Schnappinger, Dirk, Sabine Ehrt, Martin I. Voskuil, Yang Liu, Joseph A. Mangan, Irene M. Monahan, Gregory Dolganov, et al. 2003. “Transcriptional Adaptation of Mycobacterium Tuberculosis within Macrophages: Insights into the Phagosomal Environment.” Journal of Experimental Medicine 198 (5): 693–704. 10.1084/jem.20030846.

Sciara, Giuliano, Steven G. Kendrew, Adriana E. Miele, Neil G. Marsh, Luca Federici, Francesco Malatesta, Giuliana Schimperna, Carmelinda Savino, and Beatrice Vallone. 2003. “The Structure of ActVA-Orf6, a Novel Type of Monooxygenase Involved in Actinorhodin Biosynthesis.” EMBO Journal 22 (2): 205–15. 10.1093/emboj/cdg031.

Šebest, Peter, Marie Brázdová, Miroslav Fojta, and Hana Pivo. 2015. “Differential Salt-Induced Dissociation of the P53 Protein Complexes with Circular and Linear Plasmid DNA Substrates Suggest Involvement of a Sliding Mechanism.” International Journal of Molecular Sciences 16: 3163–77. 10.3390/ijms16023163.

Sun, Xian, Lu Zhang, Jun Jiang, Mark Ng, Zhenling Cui, Juntao Mai, Sang Kyun Ahn, et al. 2018. “Transcription Factors Rv0081 and Rv3334 Connect the Early and the Enduring Hypoxic Response of Mycobacterium Tuberculosis.” Virulence 9 (1): 1468–82. 10.1080/21505594.2018.1514237.

Suvorova, Inna A, Yuri D Korostelev, and Mikhail S Gelfand. 2015. “GntR Family of Bacterial Transcription Factors and Their DNA Binding Motifs: Structure, Positioning and Co-Evolution.” PLoS ONE 10 (7): 1–21. 10.1371/journal.pone.0132618.

Vigouroux, Armelle, Thibault Meyer, Anaïs Naretto, Pierre Legrand, Magali Aumont-Nicaise, Aurélie Di Cicco, Sébastien Renoud, et al. 2021. “Characterization of the First Tetrameric Transcription Factor of the GntR Superfamily with Allosteric Regulation from the Bacterial Pathogen Agrobacterium Fabrum.” Nucleic Acids Research 49 (1): 529–46. 10.1093/nar/gkaa1181.

Vindal, Vaibhav, Sarita Ranjan, and Akash Ranjan. 2007. “In Silico Analysis and Characterization of GntR Family of Regulators from Mycobacterium Tuberculosis.” Tuberculosis 87 (3): 242–47. 10.1016/j.tube.2006.11.002.

Wriggers, Willy, Sugoto Chakravarty, and Patricia A Jennings. 2005. “Control of Protein Functional Dynamics by Peptide Linkers.” Biopolymers 80 (6): 736–46. 10.1002/bip.20291.

Zeng, Jie, Wanyan Deng, Wenmin Yang, Hongping Luo, Xiangke Duan, Longxiang Xie, Ping Li, et al. 2016. “Mycobacterium Tuberculosis Rv1152 Is a Novel GntR Family Transcriptional Regulator Involved in Intrinsic Vancomycin Resistance and Is a Potential Vancomycin Adjuvant Target.” Scientific Reports 6 (May): 1–12. 10.1038/srep28002.

